# Probing the decision-making mechanisms underlying choice between drug and nondrug rewards in rats

**DOI:** 10.1101/2020.11.14.382622

**Authors:** Youna Vandaele, Magalie Lenoir, Caroline Vouillac-Mendoza, Karine Guillem, S.H. Ahmed

**Affiliations:** Lausanne University Hospital, Department of Psychiatry, 1008 Prilly, Switzerland; Université de Bordeaux, Institut des Maladies Neurodégénératives, UMR 5293, 146 rue Leo Saignat, F-33000 Bordeaux, France; CNRS, Institut des Maladies Neurodégénératives, UMR 5293, 146 rue Leo Saignat, F-33000 Bordeaux, France

**Keywords:** Choice, decision-making, sequential choice model, behavioral ecology, deliberation, habit, goal-directed, cocaine, saccharin, rats

## Abstract

Investigating the decision-making mechanisms underlying choice between drug and nondrug rewards is essential to understand how their alterations can contribute to substance use disorders. However, despite some recent effort, this investigation remains a challenge in a drug choice setting, notably when it comes to delineate the role of goal-directed versus habitual control mechanisms. The goal of this study was to try probing these different mechanisms by comparing response latencies measured during sampling (i.e., only one option is available) and choice trials. A deliberative goal-directed control mechanism predicts a lengthening of latencies during choice whereas a habitual control mechanism predicts no change in latencies. Alternatively, a race-like response competition mechanism, such as that postulated by the behavioral ecology-inspired Sequential Choice Model (SCM), predicts instead a shortening of response latencies during choice compared to sampling. Here we tested the predictions of these different mechanisms by conducting a systematic retrospective analysis of all cocaine versus saccharin choice experiments conducted in rats in our laboratory over the past 12 years. Overall, we found that rats engage a deliberative goal-directed mechanism after limited training, but shift to a SCM-like response selection mechanism after more extended training. The latter finding suggests that habitual control is engaged in a choice setting via a race-like response competition mechanism, and thus, that the SCM is not a general model of choice, as formulated initially, but a specific model of habitual choice.

## Introduction

Investigating the decision-making mechanisms underlying choice between drug and nondrug rewards is essential to understand how their alterations can contribute to substance use disorders. Over the past decade, a growing research effort has been made to model this choice situation in animals, particularly in rodents [1–3]. Overall, when given a choice, most rats prefer the nondrug alternative (e.g., sweet water; social interaction) over potent drugs of abuse, such as cocaine or heroin [4–10]. Only few individual rats prefer the drug. This pattern of individual preferences is observed even after extended drug use and regardless of the drug dose available [4, 6]. Recently, we found that preference for the nondrug option in the majority of rats is under habitual control, not goal-directed control, as it persists even after its devaluation [11, 12]. This was observed after extended training, but we do not know if habitual control of preference is due to extended training or other factors known to promote habitual control, such as, for instance, prior drug exposure [13–17]. In addition, we currently ignore if preference for the drug is also under habitual control, mainly because there is no effective method of drug reward devaluation in animals, particularly for drugs administered intravenously [18, 19]. An alternative approach, sufficiently versatile to probe the decision-making mechanisms underlying individual preferences in a drug choice setting, is thus needed.

Numerous studies suggest that there is a speed-accuracy tradeoff in the exercise of goal-directed and habitual decision-making processes [20–22]. Indeed, it is generally assumed that goal-directed behaviors are flexible but require higher cognitive demand and are time-consuming whereas habitual behaviors are inflexible in adapting to new environmental contingencies but are faster to execute. Based on this assumption, the decision-making mechanisms underlying choice should be, in theory, inferable from a detailed analysis of response latencies. Specifically, if choice behavior is goal-directed and involves a deliberation over the values of the different reward options, one would expect an increase in response latency during choice trials in comparison to sampling trials where the different options are presented separately and successively. In contrast, if behavior is habitual and performed automatically without representation of the values of the options, response latencies during choice and sampling trials should be similar. Furthermore, since behavior typically transitions from goal-directed to habitual control after extended training [23–25], the predicted difference in response latency between choice trials and sampling trials should be observed after limited training but not after extended training. Thus, analysis of response latencies has the potential to help us distinguishing between goal-directed and habitual control of preference.

A similar approach was previously used to assess the validity of different models of animal choice, including behavioral ecology-inspired models [26–29]. One such model, the Sequential Choice Model (SCM), is of particular interest here because it makes a prediction that is different from both goal-directed and habitual control models (see below). This model explains choice without postulating any explicit deliberation and/or valuation processes. Specifically, when facing different options, animals would consider each option sequentially, not simultaneously, and would decide whether to accept or reject it with no consideration of the other option available. It is hypothesized that such sequential decision process has evolved as an adaptation to the natural “reward ecology” of most animals. Unlike in laboratory choice settings, successive encounters with different rewards are the rule in natural environments while simultaneous encounters are the exception [30, 31]. Based on this assumption, the SCM proposes that there would be no genuine decision and that preference would be the result of a race-like competition between independent responses. Briefly, during choice, each option would automatically elicit a specific response with a certain latency: the shorter the latency, the more likely the corresponding response will win the race and thus, the more likely the animal will prefer that option. Importantly, due to this race-like selection process, the SCM uniquely predicts that the choice latencies should be shorter, not longer, than the sampling latencies and this, independently of the degree of prior training (see the Supplement for additional information).

Thus, different models make different testable predictions about the differences between choice and sampling latencies (i.e., longer, similar or shorter) (Table 1). Interestingly, however, these different models also make one general common prediction regarding the relationship between sampling latencies and preference, though for different reasons. All models predict that individuals that respond faster for one option relative to its alternative during sampling trials should also choose the former more frequently than the later during choice trials. In other terms, there should be a positive correlation between individual sampling latencies and individual preferences during choice. According to the SCM, the nature of this relationship should also be predictive since response latencies play a causal role in the establishment of preference in this model [28, 29]. In the other two models, sampling latencies rather represent another measure of the options relative values.

**Table 1:** Unique predictions of the different decision-making models.

| Models: | Predictions: |
| --- | --- |
| Goal-directed decision-making | Sampling latencies < Choice latencies |
| Habitual decision-making | Sampling latencies = Choice latencies |
| Sequential choice model | Sampling latencies > Choice latencies |

The goal of the present study is to test these various predictions by conducting a systematic retrospective analysis of all cocaine versus saccharin choice experiments that have been conducted in rats in our laboratory over the past 12 years. This allowed us to probe the decision-making mechanisms underlying choice between drug and nondrug rewards as a function of prior training (limited or extended).

## Results

### Lengthening of choice latencies after limited training

We first assessed distributions of response latencies and preference in experiments without prior instrumental training before choice testing (Table 2, W/O training set), assuming that behavior would be goal-directed under these conditions. All three decision-making models predict that the relative latency of each response option when encountered sequentially (i.e., sampling phase) should predict which one will be selected when encountered simultaneously (i.e., choice phase). That is, the fastest the response for an option during sampling, the more likely this option will be selected during choice. We first tested this prediction by correlating the relative response latency during sampling trials with the preference. We found that both LR and WL were positively correlated with the percentage of cocaine choice (Fig 1A-B; LR: R=0.62, p<0.001; WL: R=0.42, p<0.05). These results suggest that the faster animals respond for cocaine relative to saccharin during sampling trial, the more they would prefer cocaine.

**Table 2:** Summary and conditions of experiments included in the analysis.

| N°Exp | Exp set | N | Prior training | Training session limit | Nb training sessions | Other conditions | Nb choice sessions (FR2) | Selected choice sessions |
| --- | --- | --- | --- | --- | --- | --- | --- | --- |
| 1 | W/O training | 9 | N/A | N/A | 0 | *** | 5 | s13-s15 |
| 2 | W/O training | 11 | N/A | N/A | 0 | *** | 5 | s13-s15 |
| 3 | W/O training | 11 | N/A | N/A | 0 | *** | 5 | s13-s15 |
|  | Total : | 31 |  |  |  |  |  |  |
| 4 | W/ training | 18 | FR1 | 30 rewards/3h | 26 | *** | 8 | s32-s34 |
| 5 | W/ training | 19 | FR1 | 30 rewards/3h | 26 | *** | 8 | s32-s34 |
| 6 | W/ training | 11 | FR1 | 30 rewards/3h | 18 | 1-month HC saccharin access | 10 | s26-s28 |
| 7 | W/ training | 22 | FR1/FR2 | 30 rewards/3h | 26 | cannula intra-OFC | 10 | s34-s36 |
| 8 | W/ training | 21 | FR1 | 30 rewards/3h | 23 | cannula intra-OFC | 9 | s30-s32 |
| 9 | W/ training | 12 | FR1/FR2 | 30 rewards/3h | 21 | *** | 9 | s28-s30 |
| 10 | W/ training | 23 | FR1/FR2 | 30 rewards/3h | 32 | *** | 5 | s35-s37 |
| 11 | W/ training | 9 | FR1 | 30 rewards/3h | 19 | *** | 5 | s22-s24 |
| 12 | W/ training | 15 | FR1/FR2 | 20 rewards/2h | 21 | *** | 5 | s24-s26 |
|  | Total : | 150 |  |  |  |  |  |  |

**Figure 1:**
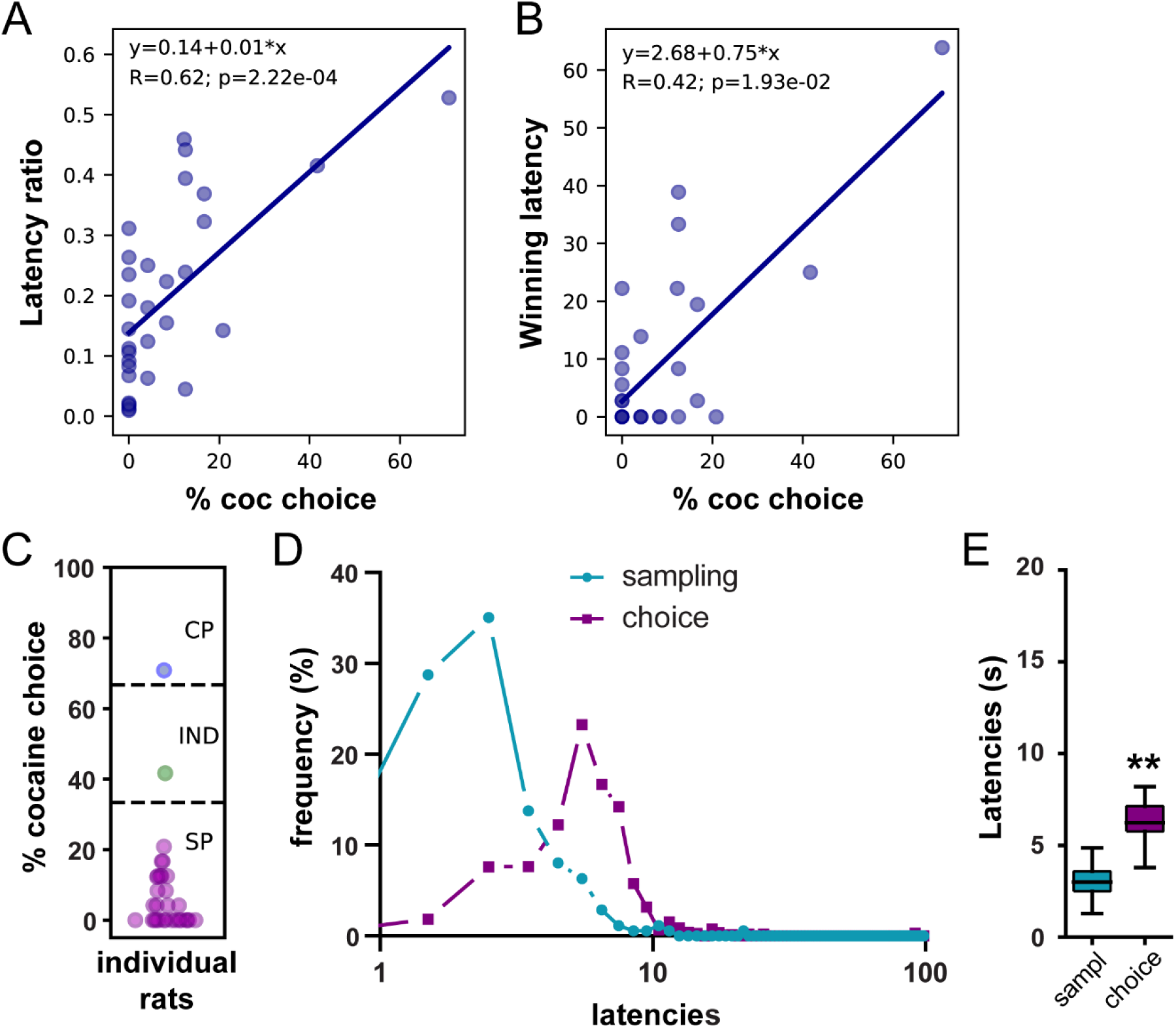
Lengthening of saccharin choice latencies compared to saccharin sampling latencies in the W/O training set. (A) Correlation between the latency ratio and the percentage of cocaine choice. (B) Correlation between the winning latency and the percentage of cocaine choice. (C) Distribution of preference scores. Only SP rats (purple circles; N=29) were considered in the analysis. (D) Distributions of sampling and choice latencies during saccharin trials in SP rats. (E) Box plot of saccharin sampling and choice latencies. The box extends from the lower to the upper quartile values with a horizontal line at the median. The whiskers extend from the box at 1.5 times the interquartile range. ** p<0.0001.

To determine whether a lengthening of choice latencies could be observed when choice behavior is presumably goal-directed, we analyzed the distribution of sampling and choice latencies during saccharin trials in saccharin-preferring (SP) rats. Among the 31 rats included in the analysis, 29 expressed a preference for saccharin (Fig 1C). In this subgroup, the distribution of choice latencies was shifted to the right compared to the distribution of sampling latencies (Fig 1D). Accordingly, choice latencies were significantly longer than sampling latencies (Fig 1E; N=29, Wilcoxon-test: p<0.0001; effect size: −0.99). These results are consistent with the involvement of a deliberative and goal-directed decision-making mechanism.

### Shortening of choice latencies after extended training

We next assessed preference and response latencies during sampling and choice in experiments including prior instrumental training before choice testing, assuming that at least responding for saccharin would be under habitual control, based on prior studies [11, 12]. We first tested the general common prediction that sampling latencies should correlate with preference and found that, similarly to the W/O training set, both LR and WL were positively correlated with the percentage of cocaine choice (Fig 2A-B; LR: R=0.67, p<0.0001; WL: R=0.64, p<0.0001), indicating that fast responses for cocaine during sampling trials are associated with higher preference for this option during choice.

**Figure 2:**
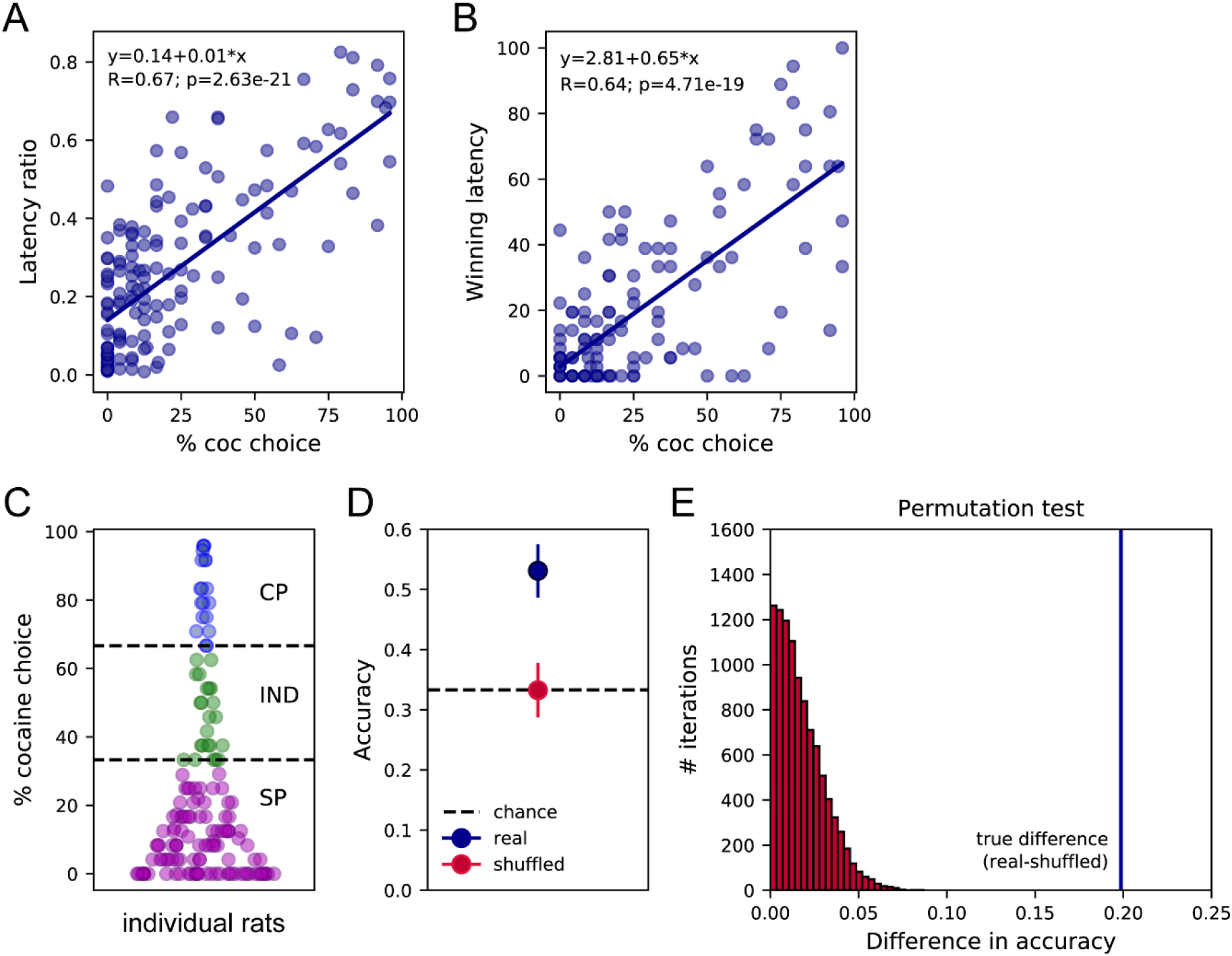
Sampling latencies correlate with preference and predict preference profiles. (A) Correlation between the latency ratio and the percentage of cocaine choice. (B) Correlation between the winning latency and the percentage of cocaine choice. (C) Distribution of preference scores and assignment of preference profiles in individual rats. SP: Saccharin-preferring rats, purple circles; IND: Indifferent rats, green circles; CP: Cocaine-preferring rats, blue circles. (D) The mean decoding accuracy (± standard deviation) of the preference profile, SP, IND or CP, of individual rats based on their mean cocaine and saccharin sampling latencies (real – dark blue) is compared with the decoding accuracy expected from chance (Shuffled – red; Chance level 33.3% – horizontal dashed line). (E) Permutation test. The true difference between accuracy scores real-shuffled (vertical blue line) is compared to the distribution of differences in accuracy following permutations with 10000 iterations. P<0.0001.

The SCM not only predicts that preference should be correlated with the relative sampling speed but also proposes that preference could be directly predicted from sampling latencies [28, 29]. To test this hypothesis, we trained a LDA model on the mean cocaine and saccharin sampling latencies to classify the preference of individual rats as saccharin-preferring (SP), indifferent (IND) or cocaine-preferring (CP). As expected from prior studies [4, 6], the majority of rats preferred saccharin with a preference score below 33.3% (SP rats: N=109, 72.7%; Fig 2C). A subset of rats were considered as indifferent with preference scores ranging between 33.3 and 66.6% (IND rats: N=23, 15.2%) or preferred cocaine with more than 66.6% of cocaine choice (CP rats: N=19, 12.6%; Fig 2C). It was possible to predict the preference profile (SP, CP or IND) from the mean cocaine and saccharin sampling latencies with an accuracy of 53.1±4.4% (Fig 2D). A permutation test confirmed that the decoding accuracy significantly departed from chance, assessed by shuffling the preference profile (Fig 2D-E). This result further validates the first SCM prediction by showing that the preference profile can be predicted above chance, from the options’ sampling latencies.

Goal-directed, habitual and sequential choice models make different predictions when comparing sampling and choice latencies (Table 1). To test these predictions, we compared the individual distributions of sampling and choice latencies for each reward, separately. To avoid any selection bias resulting from saccharin preference in the majority of rats, choice and sampling latencies distributions were compared within-subjects in different preference groups, separately.

In SP rats, the distribution of sampling and choice latencies on saccharin trials did not significantly differ, although there was a trend toward longer choice latencies (Fig 3A; N=109; Wilcoxon-test: p=0.077; effect size: −0.19). However, in CP rats, we observed a leftward shift in the distribution of latencies during cocaine choice trials compared to cocaine sampling trials (Fig 3B). Accordingly, cocaine choice latencies were significantly shorter than cocaine sampling latencies (Fig 3B; N=19, Wilcoxon-test: p<0.01; effect size: 0.75). Indifferent (IND) rats selected both cocaine and saccharin during choice trials allowing for similar ranges in the number of cocaine and saccharin choice latencies (i.e. between 8 and 16 latencies per reward). When comparing saccharin trials, we observed no difference between sampling and choice latencies, similarly to saccharin trials in SP rats (Fig 3C; N=23; Wilcoxon-test: p>0.9; effect size: −0.0072). However, during cocaine trials, the distribution of choice latencies was shifted to the left compared to the distribution of sampling latencies. Likewise, choice latencies were significantly shorter than sampling latencies (Fig 3D; N=23; Wilcoxon-test: p<0.0001; effect size: 0.96). These results partially validate the second prediction of the SCM, since choice latencies were shorter than sampling latencies for the cocaine option but not for the saccharin option. Importantly, although rare, omissions were more frequent during sampling than choice trials, which could have biased the results reported here. However, similar results were found when omission trials were excluded (saccharin trials in SP rats: p=0.06; Cocaine trials in CP rats: p<0.01; saccharin trials in IND rats: p>0.9; cocaine trials in IND rats: p<0.0001).

**Figure 3:**
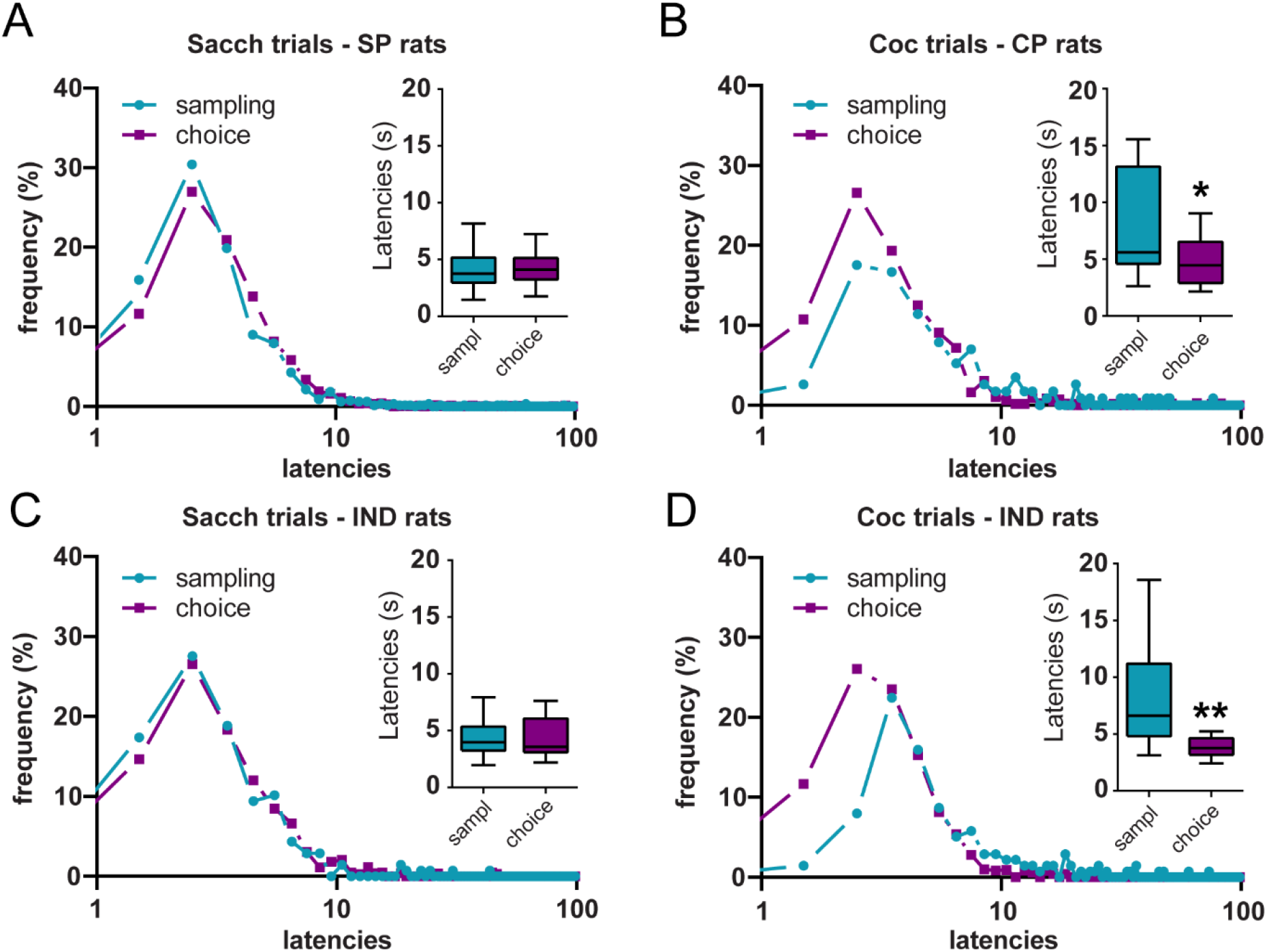
Shortening of cocaine choice latencies compared to cocaine sampling latencies. (A-D) Distributions of sampling and choice latencies during saccharin trials in SP rats (A), cocaine trials in CP rats (B) and saccharin (C) or cocaine (D) trials in IND rats. Insets: Box plots of sampling and choice latencies. Boxes extend from the lower to the upper quartile values with a horizontal line at the median. The whiskers extend from the box at 1.5 times the interquartile range. * p<0.01, ** p<0.0001.

## Discussion

The aim of this study was to determine the decision-making processes underlying choice by comparing sampling and choice response latencies, after limited or extended training. An increase in latencies during choice is predicted from the deliberative goal-directed decision-making model whereas latencies should remain the same under habitual control after extended training. Alternatively, the SCM predicts a shortening of latencies during choice compared to sampling, independently of the training duration. Here we tested these different predictions in a systematic retrospective analysis of all the choice experiments conducted in the laboratory over the past 12 years. Our analysis shows a lengthening of choice latencies after limited training, when behavior is presumably goal-directed, suggesting the involvement of a deliberative process. However, when prior training is included before choice testing, we observed similar choice and sampling latencies on saccharin trials and a shortening of choice latencies on cocaine trials. These results are consistent with both the SCM and habitual learning.

In agreement with previous experiments [27–29, 34–37], results from our analysis validated in both experimental sets (with or without prior training) the general common prediction that the relative response latency during sampling trials should correlate with preference during choice trials. Indeed, the parameters of relative latency were both strongly correlated with the percentage of cocaine choices. Furthermore, in agreement with the SCM, which assumes that the relative sampling latency causally determines preference, sampling latencies could predict whether rats preferred cocaine, saccharin or were indifferent. However, these findings are not sufficient to favor the SCM over the habitual and goal-directed choice models, which also predict (or do not exclude) a relationship between sampling latencies and preference. For instance, in a deliberative goal-directed choice model, options are compared and selected based on their relative value [26, 38, 39] and numerous studies have reported that latencies are inversely related with options’ relative value [28, 40–43]. Thus, sampling latencies and preference should be correlated under goal-directed control. More generally, it was suggested that response latencies could be used as a sensitive metric for response strength and subjective value [28, 40–43]. To our knowledge, the habitual choice model makes no specific prediction about the relationship between sampling latencies and preference, but the engagement of habit cannot be excluded based on these results. The analysis and comparison of sampling and choice latencies is therefore needed to arbitrate between the different models.

Contrary to all prior studies testing predictions of the SCM [26–29, 34–37, 44], analysis of experiments without instrumental training prior to choice testing reveals a lengthening of choice latencies compared to sampling latencies. These results are consistent with deliberative models of decision making which assume that processing of information during simultaneous encounters would increase the response latency compared to sequential encounters, due to the time cost of the evaluation process [20, 22, 26, 38, 39] (Table 1). Interestingly, in these experiments, reward-seeking behavior is presumably goal-directed. Indeed, habit typically develops across extended training [23–25], particularly when rats are trained under ratio schedule as in our experiments [45, 46]. Furthermore, in these conditions, rats were less exposed to cocaine, also known to promote habit [13–16]. The discrepancy with prior findings could result from the relatively long training in instrumental tasks involving discrete trial designs, shown to promote rapid habitual learning [47, 48] or from species differences [26–29, 34–36, 44]. Indeed, in this analysis, the lengthening of latencies was observed in naïve rats trained for a very limited number of sessions while most studies investigating SCM predictions were conducted in European starlings or pigeons (but see [37]), with an extensive instrumental training.

In the dataset with prior instrumental training, the shortening of cocaine response latencies during choice and the absence of difference for saccharin trials is consistent with both the SCM and habitual learning. We have previously shown that preference for the nondrug reward is habitual in the discrete trial choice procedure [11, 12]. In contrast to goal-directed behaviors which rely on a representation of the outcome value, habits are mediated by a stimulus-response association and do not require processing of the outcome value to be executed. Thus, under habitual control, an option should be selected with the same latency during choice and sampling (Table 1), which is what we observed for saccharin trials, in SP and IND rats. However, if rats select options according to the SCM, a shortening of choice latencies should be observed. Indeed, the race-like competition process implies that only the shortest latencies should be expressed during choice trials, the selection of one option excluding the opportunity to select the alternative option (Supplemental Figure 1). This cross-censorship between distributions of latencies results in a shortening of choice latencies. This prediction was validated for cocaine but not saccharin trials. The negative result for saccharin trials is however consistent with prior studies testing SCM predictions [27–29, 35–37, 44, 49]. Indeed, none of these studies but one [37] has reported a shortening of choice latencies for the preferred option. Demonstrating this decrease in latency is difficult because when animals strongly prefer one option, the distribution of latencies for that option is minimally censored (Supplemental Figure 1). Furthermore, the high relative value of this option results in short latencies during sampling, limiting the room for further decrease. Analysis of prior studies conducted in our laboratory indicates that response latencies cannot go below 3-s, likely representing the reaction time.

Although cocaine was the preferred option in CP rats, we were able to observe a shortening of latencies at choice. In contrast to saccharin, cocaine is generally selected with longer latencies during sampling, preventing the floor effect. Longer latencies to select cocaine could result from the ambivalent and anxiogenic effects of the drug [50–53] or from its delayed pharmacological effects on the brain [54]. Interestingly the gap between cocaine choice and sampling latencies was more pronounced in IND rats compared to CP rats. This result further supports the SCM, which predicts that trimming of the leftward side of the distribution of latencies would be more pronounced when the overlap between response distributions is stronger [26, 28] (Supplemental Figure 1). Since rats are indifferent, the drug and nondrug rewards are closer in value and selected with more comparable response latencies during sampling. The stronger cross-censorship between distributions of latencies during choice could have increased the shortening of cocaine choice latencies in this subgroup of rats. However, at indifference, the SCM predicts a shortening of choice latencies for both options [44]. The similarity between saccharin sampling and choice latencies in IND rats is thus unexpected. Given the already low value of latencies during sampling (mean: 4.7±0.49s; median: 3.97s), a floor effect cannot be excluded.

Like habit, the automatic selection process of option based on the response latency implied by the SCM, is oblivious to the options’ value. The SCM assumes that during sequential encounters, each option are processed independently and choice results from a race between the mechanisms generating response latencies for each available option [26]. Thus, although the SCM proposes a mechanism for the selection of responses based on their latency, this model makes no assumption on the valuation process generating distributions of latencies. It is tempting to speculate that the habitual system would process the action value (or “action policy”) while the SCM offers a mechanism by which habit translates in preference in a choice setting between two simultaneous options. This assumption is in agreement with a recent theory suggesting that decision-making does not necessarily require computation of the economic value by the brain [55]. Overall, our analysis suggests that habitual responding for the drug and the nondrug reward is engaged after extended training and selected via a race-like competition mechanism, thus indicating that the SCM is not a general model of choice as initially proposed [26, 28], but rather constitutes a specific model of habitual choice.

To conclude, this systematic analysis has begun to probe the decision-making processes underlying choice between sweet water and cocaine in rats. It shows that when simultaneously facing these two options, rats first engage a deliberative strategy with computation and comparison of the options value. However, after more extended training, rats adopt a SCM-like response selection process while their behavior becomes habitual. Clearly, more research is needed to determine the time-course and conditions of habitual learning for the drug and nondrug rewards in our discrete-trial choice procedure. However, these results suggest that the comparison of sampling and choice latencies could be used as a proxy to determine whether behavior is habitual or goal-directed. Given the difficulty to devalue drugs administered intravenously, such analysis of latencies could be particularly useful to infer the nature of behavioral control in drug settings. Furthermore, this study offers a new framework to understand how habit can translate in preference in a choice setting between a drug and a nondrug reward. This theoretical framework will ultimately allow progress in our understanding of the relation between habits and choice and their respective role in substance use disorder [56].

## Materiel and methods

### Subjects

The data analyzed in this study have been obtained from previous experiments conducted in our laboratory over the past 12 years. All experimental subjects were male adult Wistar rats (Charles River, L’Arbresle, France). Rats were housed in groups of two or three and maintained in a temperature-controlled vivarium (23° C) with a 12-h reverse light-dark cycle. Testing occurred during the dark phase of the cycle and water and food were available ad libitum in all experiments. All experiments were carried out in accordance with institutional and international standards of care and use of laboratory animals [UK Animals (Scientific Procedures) Act, 1986; and associated guidelines; the European Communities Council Directive (2010/63/UE, 22 September 2010) and the French Directives concerning the use of laboratory animals (décret 2013-118, 1 February 2013)]. All experiments have been approved by the Committee of the Veterinary Services Gironde, agreement number B33-063-5.

### Initial operant training

In the first set of experiments, a total of 31 rats were directly tested in the choice schedule without prior operant training (Table 2; W/O training set).

In the second set of experiments, a total of 150 rats were first trained several weeks (3-5) under a fixed-ratio (FR) schedule of saccharin and cocaine self-administration on alternate daily sessions, six days a week, before choice testing (Table 2; W/ training set). FR training allowed rats to learn the value of each reward and to associate its delivery with a different response option before choice testing. On saccharin sessions, lever pressing on the saccharin lever was rewarded by a 20-s access to water sweetened with 0.2% of sodium saccharin delivered in the adjacent drinking cup. During the first 3-s of each 20-s access to sweet water, the drinking cup was filled automatically with sweet water; during the next 17-s, additional volumes of sweet water were obtained on demand by voluntary licking. On cocaine sessions, lever pressing on the alternative lever was rewarded by one intravenous dose of cocaine (0.25 mg delivered over 4 seconds). For both cocaine and saccharin sessions, reward delivery initiated a concomitant 20-s time-out period signaled by the illumination of the cue-light above the available lever. During the time-out period, responding had no scheduled consequences. Sessions ended after rats had earned a maximum of 20-30 saccharin or cocaine rewards or 2-3 h had elapsed (Table 2).

### Discrete-trials choice protocol

Each daily choice session consisted of 12 trials, spaced by 10min inter-trials intervals, and divided into two successive phases, sampling and choice. The sampling phase was composed of 4 sampling trials in which each lever and thus each response option was presented alternatively and sequentially. If rats responded twice within 5 min on the available lever, they were rewarded by the corresponding reward. Reward delivery was signaled by the immediate retraction of the lever and illumination of a cue-light above it. If rats failed to complete the response requirement within 5 min, the lever was retracted until the next trial 10 min later. The choice phase consisted of eight choice trials during which the two response options were presented simultaneously. Specifically, each choice trial began with the simultaneous presentation of both levers S and C and rats could select one of the two levers by responding twice consecutively on it to obtain the corresponding reward. Reward delivery was signaled by the simultaneous retraction of both levers and illumination of the cue-light above the selected lever. If rats failed to respond on either lever within 5 min, both levers were retracted and no cue-light and reward was delivered. A schematic diagram of the forced choice trials procedure can be found in [32, 33].

In the W/O training set, rats were first trained in the discrete-trials choice schedule with a FR1 for 10 sessions before testing with the final FR2 schedule for 5 sessions.

### Systematic analysis

#### Selection of experiments and data analysis

Only choice experiments that were conducted under similar conditions (e.g., initial FR training, conditions of reward delivery, inter-trial intervals, …etc.) and that resulted in a stable preference within 5-10 choice sessions (i.e., no increasing or decreasing trend and significant correlation between preference scores over the last 3 sessions) were included in the present analysis (Table 2).

Sampling and choice latencies for each response option were analyzed for each individual rat over the last three stable choice sessions. In total, there were 6 sampling latencies per option and per rat and 24 choice latencies, with a variable number of responses for cocaine and saccharin, depending on the rat’s preference. For convenience, performance during choice was expressed in percent of cocaine choices. Response latencies corresponded to the time to complete the FR2 requirement from trial onset (signaled by the lever insertion). When a rat failed to respond within 5 min after trial onset (omission), it was assigned a maximal response latency of 300 seconds. However, omissions occur rarely.

#### Test of the general common prediction

To estimate the relative speed at which the cocaine and saccharin options are selected during sampling trials, we computed for each individual rat the latency ratio (LR) and the proportion of winning latencies (WL). The LR was computed by dividing the mean saccharin sampling latency by the sum of saccharin and cocaine mean latencies. Thus, LR values close to zero indicate faster saccharin sampling latencies while LR values close to one indicate faster cocaine sampling latencies. The WL was computed by estimating the probability that each of the six cocaine sampling latencies was shorter (i.e., a win) than each of the six saccharin sampling latencies when compared two by two. The WL values ranged from 0 (all cocaine latencies are longer than the longest saccharin latency or cocaine never wins the race) to 100% (all cocaine latencies are shorter than the shortest saccharin latency or cocaine always wins the race). Correlation analyses were conducted between these two measures of relative latencies and the percentage of cocaine choices.

To determine whether and to what extent, cocaine and saccharin sampling latencies could predict rat’s preference profile, a linear discriminant analysis (LDA) model (LinearDiscriminantAnalysis from sklearn library in Python) was trained on the mean cocaine and saccharin sampling latencies to classify the preference of individual rats as saccharin-preferring (SP), indifferent (IND) or cocaine-preferring (CP). Rats were considered as SP, IND or CP if their preference was below 33.3%, between 33.3-66.6% or above 66.6%, respectively. LDA models were trained on 90% of the dataset and used to classify rats’ preference in the remaining 10% (stratified 10-Fold cross-validation with 20 iterations; RepeatedStratifiedKFold from sklearn library in Python). To account for the unbalanced number of SP, IND and CP rats, we performed the analysis on the same number of rats in each preference group by randomly sampling subjects in the SP and IND groups based on the number of CP rats (N=19). This analysis was performed on 50 random selections of SP and IND rats and the performance across all 50 repetitions was averaged to determine the model accuracy. The same analysis was conducted with the preference group identities shuffled to determine accuracy expected by chance. To assess whether decoding accuracy significantly departed from chance, a permutation test was conducted.

#### Comparison of choice and sampling latencies

To test predictions of the different decision-making models, we compared the distributions of sampling and choice latencies for each response option. Note that the number of choice latencies (i.e. max. 24) was larger than the number of sampling latencies (6 per option) and that among choice trials, the number of saccharin responses was disproportionally higher than cocaine responses because of preference for saccharin in the majority of rats. To avoid any selection bias related to differences in preference, choice and sampling responses for saccharin were compared in SP (<33.3% cocaine choice) and IND rats (33.3-66.6% cocaine choice) whereas choice and sampling responses for cocaine were compared in CP (>66.6% cocaine choice) and IND rats. Note that the IND group comprises comparable numbers of cocaine and saccharin choice latencies allowing for analysis of both reward responses in this subgroup of rats.

#### Statistical analysis

Linear regressions were tested with the Spearman’s rank correlation test. Mean sampling and choice latencies were compared within subject using the non-parametric Wilcoxon test. The effect sizes were estimated with the rank-biserial correlation. Following the linear discriminant analysis, a permutation test was conducted to determine whether the decoding accuracy significantly departed from chance. All statistical analyses were conducted on Python.

## Authors contributions

Conceptualization: YV, SHA; Formal analysis: YV; Investigation: ML, KG, CVM; Visualization: YV; Resource: SHA; Writing - Original Draft Preparation: YV; Writing – Final Version: YV, SHA; Writing – Review, Editing and Validation of the Final Version: YV, SHA, ML, KG, CVM

## Supplementary figure

**Supplemental figure 1:**
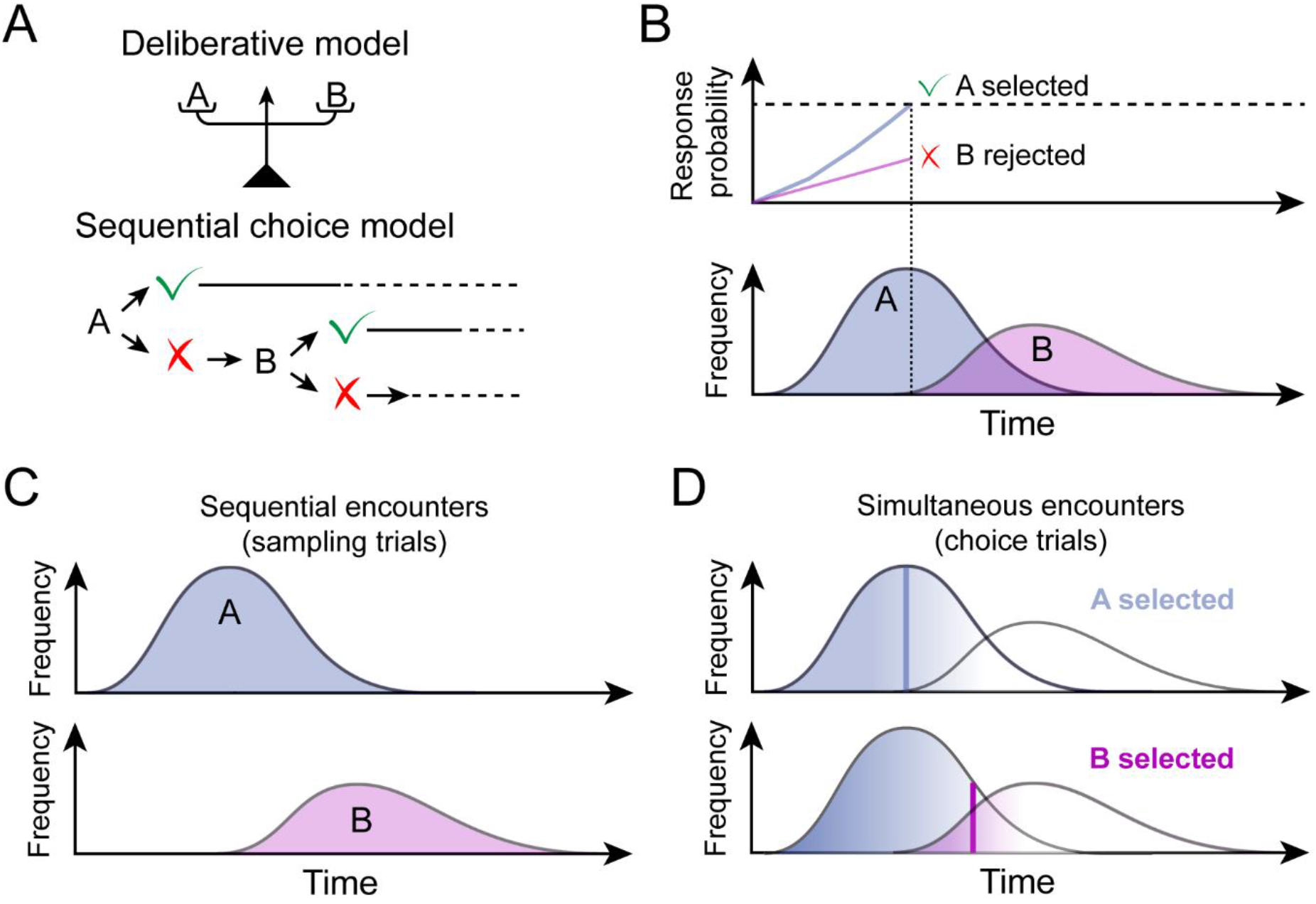
Illustration of assumptions and predictions of the sequential choice model (SCM). (A) In a deliberative decision-making model, individuals assign values to the different options and compare these values at the time of choice to select to most profitable option (top panel). Alternatively, the SCM assumes that individuals make no comparison between options to make a choice (bottom panel). Instead, decision-making mechanisms are adapted to sequential encounters, in which individuals decide to accept or reject single opportunities. (B) The SCM proposes that individuals assign independently to each option a subjective value based on the option’s profitability relative to background opportunities. This value is expressed as a probability to accept each option instead of pursuing the search (i.e. the response latency). During a choice between two options A and B, the tendency to respond for both options is compared to a threshold. The response reaching the threshold first wins the race and is selected, while the alternative response is aborted (top panel). This results in a cross-censorship between distributions of response latencies (bottom panel); only the fastest response is selected and produces a latency observation. (C) When options are presented sequentially during sampling trials, there is no cross-censorship and the distributions of latencies are entirely expressed. (D) However, when options are presented simultaneously during choice trials, the cross-censorship described in (B) leads to a shortening of latencies during choice compared to sampling trials; the preferred option A is more often selected because response latencies for this option are overall shorter (top panel) but because of the overlap in distributions of latencies, the least preferred option B can occasionally be selected (bottom panel). Choices of option A are less censored by longer responses for option B (blue gradient, top panel); thus, the expected shift toward shorter latencies is weaker for the preferred option. In contrast, choices of the least preferred option B are largely censored by responses for option A (pink gradient, bottom panel). Thus, the expected shortening of choice latencies is stronger for the least preferred option.

## Notes

### Competing Interest Statement

The authors have declared no competing interest.

